# Heterogeneous genomic architecture of skeletal armour traits in sticklebacks

**DOI:** 10.1101/2023.05.28.542672

**Authors:** Xueling Yi, Petri Kemppainen, Kerry Reid, Ying Chen, Pasi Rastas, Antoine Fraimout, Juha Merilae

## Abstract

Whether populations adapt to similar selection pressures using the same underlying genetic variants depends on population history and the distribution of standing genetic variation at the metapopulation level. Studies of sticklebacks provide a case in point: when colonising and adapting to freshwater habitats, three-spined sticklebacks (*Gasterosteus aculeatus*; TSSs) with high gene flow tend to fix the same adaptive alleles in the same major loci, whereas nine-spined sticklebacks (*Pungitius pungitius*; NPSs) with limited gene flow tend to utilize a more heterogeneous set of loci. In accordance with this, we report results of quantitative trait locus (QTL) analyses using a F_2_ back-cross design showing that lateral plate number variation in the western European lineage of NPSs mapped to three moderate-effect QTL, contrary to one major QTL in TSSs and these QTL were different from the four previously identified QTL in the eastern European lineage of NPSs. Furthermore, several QTL were identified associated with variation in lateral plate size, and three moderate-effect QTL with body size. Together, these findings indicate that genetic underpinnings of skeletal armour variation in *Pungitius* sticklebacks are more polygenic and heterogenous than those in three-spined sticklebacks, indicating limited genetic parallelism underlying armour trait evolution in the family Gasterostidae.

## Introduction

Why and when evolution can be repeatable and predictable remains a longstanding question in biology. Parallel evolution has been the prevailing hypothesis explaining repeatable phenotypic evolution (Martin and Orgogozo, 2013) by a set of identical-by-descent genetic variants controlling the same adaptive traits in multiple independent evolutionary lineages (Schluter et al. 2004). The probability of parallel evolution depends on many factors (MacPherson and Nuismer 2017; Bolnick et al. 2018), including how standing genetic variation is distributed at the metapopulation level (Fang. et al. 2020; Fang et al. 2021; Kemppainen et al. 2021; Schlötterer 2023). When populations are well connected, adaptive alleles are easily transported from one population to another and thereby available to be picked by selection in multiple populations subject to similar environmental pressures (Ralph and Coop 2015; Bailey et al. 2017; Lee and Coop 2017; Roberts-Kingman et al. 2021). However, once gene flow is sufficiently restricted, the potentially adaptive alleles may not reach all populations, forcing selection to work with alternative alleles in different loci (Merilä 2014; Kemppainen et al. 2021). The likelihood of parallel evolution is also influenced by demographic history of populations (MacPherson and Nuismer 2017; Thompson et al. 2019): parallel evolution is less likely in small populations where bottlenecks and strong genetic drift result in the stochastic loss of adaptive alleles (Leinonen et al. 2012; Fang et al. 2020; Dahms et al. 2022). Accordingly, smaller and less connected populations tend to have more heterogeneous genetic architectures underlying the same morphological transitions even if driven by similar ecological pressures, resulting in non-parallel evolutionary responses to selection. Understanding the relative contributions of parallel and non-parallel genetic changes to repeated phenotypic evolution in the wild can improve theoretical understanding of the role of natural selection in shaping genetic variation (Johanneson 2001) and inform predictions on the evolutionary potential of species and populations to cope with ongoing environmental changes (Bay et al. 2017).

While early research found ample evidence of genetic parallelism at both species and population levels (Colosimo et al. 2005; Stewart and Thompson 2009; Bernatchez et al. 2010; Elmer et al. 2010), it is now becoming increasingly clear that parallelism is perhaps not as common as previously thought (e.g., Bolnick et al. 2018; Fang et al. 2020; Kemppainen et al. 2021; Schlötterer 2023). The prevailing hypothesis of parallel evolution could thus stem from an ascertainment bias driven by the novelty value of discovering evidence for genetic parallelism, as well as from focus on systems characterized by high genetic connectivity. On the other hand, non-parallelism is more likely to dominate in species characterized by low genetic connectivity and small effective population sizes. The stickleback fishes have been popular model systems to study the repeatability of evolution during their numerous and recurrent colonisations from marine to freshwater habitats throughout past glacial cycles (e.g., Colosimo et al. 2005; Coyle et al. 2007; Marchinko and Schluter, 2007; Chan et al. 2010; DeFaveri et al. 2011, 2013; Jones et al. 2012; Merilä 2013; Fang et al. 2021; Kemppainen et al. 2021; Reid et al. 2021; Roberts-Kingman et al., 2021). What has transpired from these studies is that in the case of the three-spined stickleback (*Gasterosteus aculeatus*), adaptation to the freshwater environment is typically achieved by parallel genetic changes in major-effect loci (Colosimo et al. 2005; Chan et al. 2010; Peichel and Marques, 2017; Reid et al. 2021; but see: DeFaveri et al. 2011; Leinonen et al. 2012; Erickson et al. 2016; Fang et al. 2021). However, in the case of the nine-spined stickleback (*Pungitius pungitius*), there is much less genetic parallelism over similar geographic distances in response to adaptation to freshwater environments (Shikano et al. 2013; Fang et al. 2021; Kemppainen et al. 2021). For example, pelvic reduction is underlied by one major-effect locus in different freshwater populations of three-spined sticklebacks (Chan et al. 2010), whereas in the case of nine-spined sticklebacks it is underlied by a range of genetic architectures from single locus to polygenic systems (Kemppainen et al. 2021).

Another textbook example of predictable and repeatable evolution is the lateral plate number reduction in freshwater three-spined sticklebacks after colonization from marine habitats (Colosimo et al. 2005). The majority of their freshwater populations worldwide have achieved lateral plate reduction via inheritance of the same ancestral allele associated with the Ectodysplasin-A gene (*Eda*; Colosimo et al., 2005; O’Brown et al. 2015). Quantitative trait locus (QTL) mapping studies have revealed that the QTL containing *Eda* gene explains 69% - 98% variation in lateral plate phenotypes (Colosimo et al. 2004; Liu et al. 2014; Glazer et al. 2015; Erickson et al. 2016; Schluter et al. 2021; see also: Loehr et al. 2012), suggesting large-effect single-locus basis for this ecologically important adaptive trait. However, the *Eda* gene was not associated with lateral plate variation in nine-spined sticklebacks from North America (Shapiro et al. 2009) or eastern Europe (Yang et al. 2016). Instead, Yang et al. (2016) mapped the genetic architecture of lateral plate variation in the eastern European lineage (EL) of nine-spined sticklebacks to four different chromosomes (linkage groups LG8, LG12, LG20, LG21), none of which included the *Eda* gene. These studies thus suggest major interspecific differences in genetic architectures of lateral plate variation within the stickleback family Gasterosteidae, as well as a more polygenic and heterogeneous genetic architecture for this trait in the genus *Pungitius*.

The main aim of this study was to address the repeatable phenotypic evolution hypothesis (Martin and Orgogozo, 2013) in the context of genetic architectures of lateral plate phenotypes in *Pungitius* sticklebacks. To test the hypothesis of parallel evolution, we mapped QTL associated with variation in lateral plate numbers utilizing a F_2_-generation backcross family between two closely related *Pungitius* species: the fully plated *P. sinensis* and the partially plated *P. pungitius* from the western European lineage (WL). The WL *P. pungitiu*s has a distinct evolutionary history from the eastern European lineage (EL; Teacher et al. 2011; Guo et al. 2019; Feng et al. 2022) and has not yet been studied regarding their armour plate genetic architecture. The repeatable evolution hypothesis predicts that lateral plate number variation in the WL should be controlled by allelic variation either in the same genetic locus as that in the sister species three-spined stickleback or the previously identified QTL in the EL *P. pungitiu*s. The alternative hypothesis of non-parallel evolution of the same morphology underlied by different QTL is also plausible in the light of previous QTL analyses of lateral plate number variation in *Pungitius* sticklebacks (Shapiro et al. 2009; Yang et al. 2016). The secondary aims of this study were to identify QTL associated with lateral plate size and body size variation in *P. pungitius,* as well as capitalize the opportunity to narrow down the location of the sex determination region in *P. sinensis* (cf. Natri et al. 2019) with the aid of a dense set of SNPs and an improved *Pungitius* reference genome (Kivikoski et al. 2021).

## Material and Methods

### Crosses and fish rearing

A fully plated female *P. sinensis* from Yoneshiro River, Odate, Japan (40°16’ N, 140°29’ E) was artificially crossed with a partially plated male *P. pungitius* from Brugse polders, Maldegem, Belgium (51°10’ N, 03°28’ E) and the resulting F_1_ hybrids were reared in a 28-l tank at 16 °C until mature. The F_1_ hybrid males of these crosses were found to be sterile (Natri et al. 2019), and thus a backcross was conducted by naturally mating a partially plated F_1_ hybrid female with an outbred fully plated *P. sinensis* male (Fig. 1). This backcross mating was carried out in a 6-l tank with a zebrafish rack system (Aquaneering Inc., USA) to obtain fertilized clutches repeatedly. From three days post-hatching, larvae were fed twice daily with newly hatched brine shrimp (*Artemia sp*.) nauplii. Approximately eight weeks post-hatching, frozen bloodworms (Chironomidae larvae) were added to their diet. The 24-h photoperiod was maintained during the rearing. Wild fish were obtained in accordance with national and institutional ethical regulations with permission from the Finnish Food Safety Authority (#1181/0527/2011 and #3701/0460/2011). All fish rearing was conducted under the license from the Finnish National Animal Experiment Board (#STH379A and #PH1236A).

**Figure 1.**
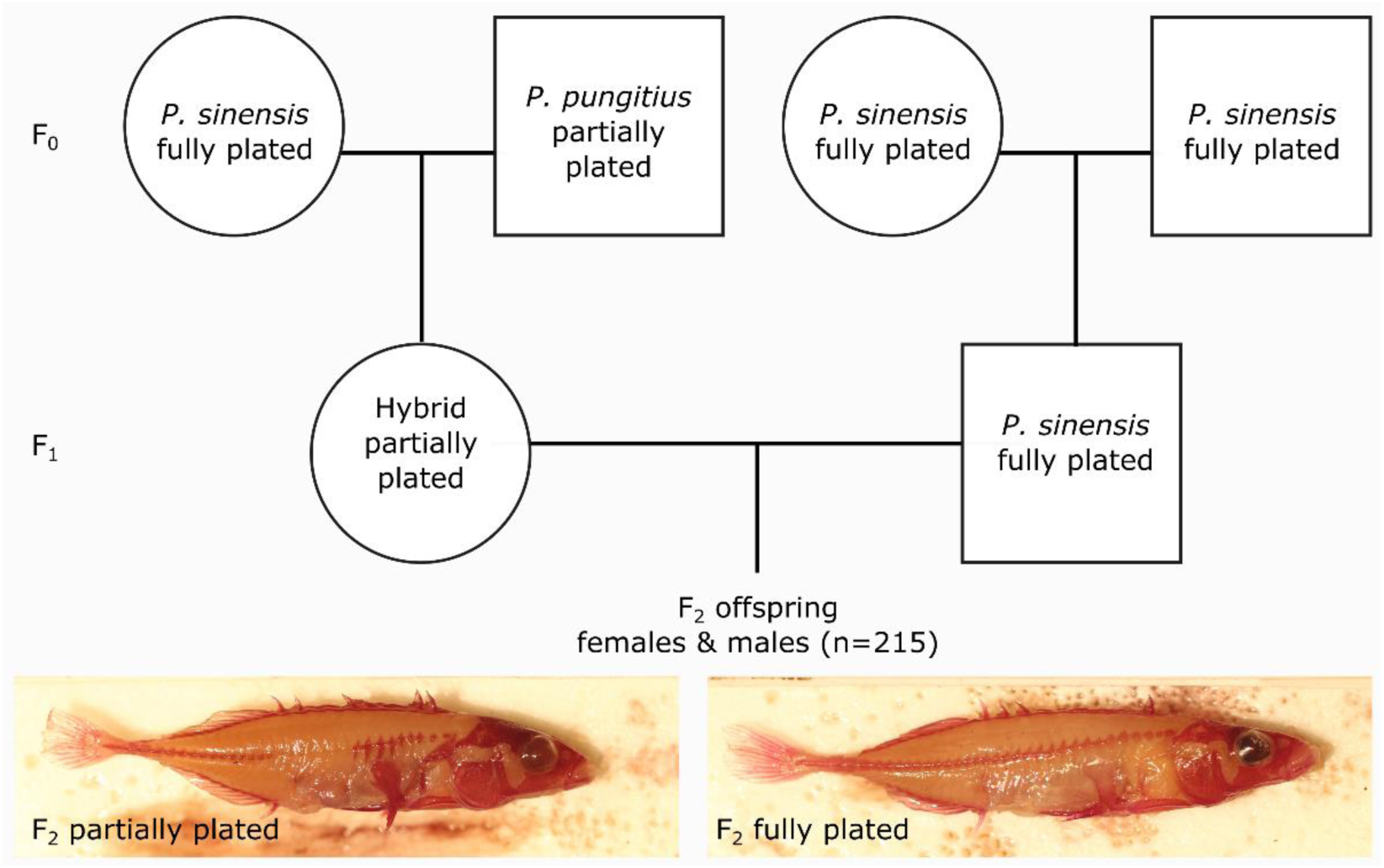
The backcross experiment between fully plated *Pungitius sinensis* from Japan and partially plated *P. pungitius* from Belgium. Circles represent females and squares represent males. The two photos show fully plated and partially plated phenotypes of the F_2_ progeny stained with alizarin red.

The F_2_ progeny were reared in 28-l tanks (20–30 fish per tank) at 16°C for 40 weeks after hatching. At the end of the experiments, the wild-caught grandparents (F_0_), F_1_ parents, and F_2_ offspring were euthanized with MS-222 (Tricaine methanesulfonate) and stored in ethanol. A fin clip was taken from all individuals and preserved at −18°C for DNA extractions. In total, 215 F_2_ offspring from six clutches were available for the analyses (Table S1). Phenotypic measurements (see below) were taken from these F_2_ offspring and their sex was recorded based on gonadal inspection after dissection.

### Phenotypic measurements

Firstly, the ethanol-stored carcasses of F_2_ offspring were fixed in 4% formalin and stained with alizarin red following the standard procedure (e.g., Trokovic et al. 2011) to visualize bony structures. Subsequently, phenotypes were quantified by ImageJ (https://imagej.nih.gov/ij/index.html) from photographs (Fig. 1; Fig. S1) of the stained fish. Standard length and maximum body height of each specimen was measured to nearest 0.01mm from the photographs.

Plate and myomere numbers were counted on both sides of the body under a dissecting microscope. The proportion of plated myomeres on each side was calculated to represent the level of platedness. The height, width and surface area of the largest lateral plates were measured on both sides of the body. Most of the largest plates were found on myomere seven counted from the anterior part of the body towards the tail (Fig. S1). If the fish had no plate in these positions, the largest plate located within the first 15 anterior myomeres were measured instead with their myomere position recorded. We did not measure plates on the myomere positions > 15 because plates in the posterior region have distinct shapes and sizes compared to those in more anterior positions. Measurements of the same phenotype on the left and right sides of the body were highly correlated (*r* > 0.86, p < 1e-10; Fig. S2). Therefore, in the following analyses we used average values of each measurement of the two sides.

The distribution of phenotypic measurements was visualized in R (R core team 2022). Traits fitting the normal distribution (namely standard length, body height, plate area, plate height, and plate width; Fig. S3) were analysed using raw values. Distributions of the plate number and the proportion of plated myomeres were left-skewed and bimodal because of the presence of only fully- and partially-plated individuals (Fig. S3). This pattern is related to our experimental design (Fig. 1) where offspring genotypes can only be *P. sinensis* homozygotes (SS, likely fully-plated) or *P. sinenesis*-*P. pungitius* heterozygotes (SP, likely partially-plated). Therefore, the missing *P. pungitius* homozygous genotypes likely account for the lack of low-plated individuals (WL *P. pungitus* have on average 5.4 lateral plates; Banarescu and Paepke 2001) and the left skew of plate number distribution. We first conducted QTL analyses on the mean number of plates (plateN_mean) as a numeric trait. To account for the lack of normality in the distribution of raw data, we also conducted QTL analyses on two transformed binary traits. First, the plate number was transformed into a binary trait, plateN_binary, where individuals were labeled as partially-plated if having the mean number of plates ≤ 25 (around the peak of the first phenotype cluster, Fig. S3) and the rest of the individuals were fully-plated. Second, the proportion of plated myomeres was transformed into the binary trait, plateness, where individuals having 100% plated myomeres on both sides were considered fully-plated while the rest were partially-plated.

### Sequencing and genotyping

DNA was extracted from the preserved fin-clips of wild-caught grandparents, F_1_ hybrid parents, and F_2_ offspring using the modified salting out method of Sunnucks and Hales (1996). Libraries for the restriction-site associated DNA sequencing (RADseq) were prepared using the *PstI* restriction enzyme (Miller et al. 2007). The prepared libraries were sequenced on an Illumina HiSeqTM 2000 (BGI Hong Kong). Demultiplexed raw sequencing data were obtained from BGI with adapter and low-quality reads removed. These reads were around 43bp in length and they were mapped to the reference genome of *Pungitius pungitius* (version 7, Kivikoski et al. 2021) using the BWA v0.7.17 backtrack algorithm (bwa aln and bwa samse; Li and Durbin 2009). The mapped reads were reformatted, sorted, and indexed using SAMtools version 1.16.1 (Danecek et al. 2021). Genotyping was conducted according to the sequencing data processing pipeline of Lep-MAP3 (Rastas 2017). Briefly, SAMtools mpileup (Li 2011) was run on the mapped data by linkage group. The version 7 reference genome has 21 linkage groups (LGs) and 1645 unassembled contigs (including a mitochondrial genome). Very few data were mapped to the unassembled contigs whose physical positions were unclear. Therefore, we only used the data mapped to the 21 linkage groups as the input of the Lep-MAP3 module Pileup2Likelihoods to estimate genotypes.

### Linkage map construction

Linkage maps were constructed following the Lep-MAP3 default pipeline (Rastas 2017). Briefly, the module ParentCall2 was run on all genotyped individuals (offspring, parents, grandparents) with the noninformative sites removed (removeNonInformative=1). The retained genotypes were processed by the module Filtering2 with the parameter dataTolerance=0.001 (i.e., segregation distortion limit 1:1000). Then the module SeparateChromosomes2 was run to assign markers to linkage groups (ordered by size) using the parameter lodLimit=50. The first (i.e., largest) 21 linkage groups were retained, corresponding to the number of chromosomes in the reference genome (Fig. S4). The linkage map was modified by collapsing all the other linkage groups with unassigned singletons. Next, the module JoinSingles2All was run using the modified linkage map and the non-filtered genotyping data (the output from ParentCall2) to map as many singletons to the 21 linkage groups as possible, using the parameters lodLimit=40, lodDifference=2, lod3, Mode=3, iterate=1, and distortionLod=1. Finally, the module OrderMarkers2 was used to order markers in the mapped 21 linkage groups and generate phased maternal maps using commands outputPhasedData=1 and informativeMask=2.

To get optimal results, OrderMarkers2 was run five times independently for each linkage group and the runs having the highest likelihoods were used as the optimal parent-phased maps. These optimal maps were then evaluated in OrderMarkers2 (command evaluateOrder) with additional parameters improveOrder=0 and grandparentPhase=1 to convert them into grandparental phases. The optimal parental phases and corresponding grandparental phases were matched using the phasematch.awk script of Lep-MAP3. The matched maps were converted back to genomic coordinates using the awk script in Lep-MAP3, and the maps were re-named by chromosomes based on the physical positions of their markers. In each map, the few markers that were located in different physical chromosomes were removed. Then the converted linkage maps were visualized by plotting the genetic position (centiMorgan, cM) against the physical position (bp), and the maps were further cleaned by manually removing the outlier markers and markers that generated gaps at the start or end of each linkage map. Finally, the map2genotypes.awk script of Lep-MAP3 was used to convert the cleaned maps into F_2_ genotypes.

### QTL mapping

The QTL mapping was conducted using the software R/qtl2 (Broman et al. 2019). The genotypes and linkage maps obtained from Lep-MAP3 were reformatted for R/qtl2 using custom scripts. Backcross (bc) cross type was specified in the control files. Pseudomarkers were inserted into genetic maps at the distance of 1 cM. Genotype probabilities were calculated using the error probability of 0.002. Genotype-phenotype associations were estimated by conducting genome scans (command scan1) using the default simple linear method of the Haley-Knott regression. The other R/qtl2 parameters were kept as default unless stated otherwise. Statistical significance thresholds were established using 1000 permutations (command scan1perm). The obtained significant threshold (p=0.05) was used to identify QTL peaks and their 95% Bayes credible intervals (CI) using the command find_peaks (peakdrop=2).

We first conducted QTL mapping on the binary phenotypic sex. Pearson’s Chi-squared test showed that phenotypic sex was not dependent on the clutch identity (p = 0.798) and thus we did not add any covariate in this model. The model was run on all 215 F_2_ offspring, including 109 phenotypic females, 105 phenotypic males, and one individual of unknown phenotypic sex. Preliminary analyses indicated misidentified phenotypic sex in 11 individuals (see supplementary information) which were removed together with the sex-unknown individual from downstream analyses for clarity.

QTL mapping on the body size measurements (standard length and body height) was carried out with sex and clutch included as additive covariates (argument addcovar) because both phenotypes showed significant dependency on sex and clutch in tests of the Analysis of Variance (ANOVA; Table S2). For the binary traits, plateN_binary and plateness, we tested their dependency on sex and clutch identities using the Pearson’s Chi-squared test and their dependency on the standard length and body height using t-tests. None of the variables showed significant relationships (*p* > 0.05) with either binary traits. Therefore, no covariate was added in the QTL mapping of plateN_binary or plateness. For each of the other numeric traits (plate number, plate area, plate height, plate width), the dependency on sex and clutch was first tested using ANOVA. Then a generalized linear model of each trait was fitted on the detected significant independent variables, and the residuals were used to further test dependency on standard length and body height (i.e., with sex and clutch controlled; Miller et al. 2014). All significant independent variables were then included as additive covariates in the QTL mapping of the corresponding trait (Table S2).

The genomic regions spanning the 95% CIs around significant QTL were considered as QTL regions. The percentage of phenotypic variation explained (PVE) by each significant QTL peak was calculated as 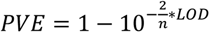 where *n* is the sample size and LOD is the logarithm of the odds of the QTL peak (Broman and Sen 2009; Smith et al. 2020). In addition, for each trait, we also used a previously described lasso regression approach to jointly incorporate all SNPs into a multilocus model to estimate the total PVE value that approximates the narrow sense heritability of a given trait (Li et al. 2017, 2018; Kemppainen et al. 2021). To facilitate calculation of the lasso algorithm, the same genotype file as used for R/qtl2 analyses was re-coded by removing duplicate SNPs that had the same genotypes (2 677 SNPs retained) and merging all SNPs outside significant QTL regions into one chromosome representing the non-QTL region. For each trait (excluding sex), the total PVE was estimated using all remaining SNPs, and the PVE of each QTL was estimated using the SNPs corresponding to the given QTL region. All phenotypic analyses and QTL mapping were conducted using R version 4 (R Core Team 2022).

### QTL effects and candidate genes

As we were mostly interested in traits related to lateral plate variation (i.e., plate number and plate size), we further estimated the effects of all the combinations of identified QTL peaks in these traits. QTL effects were estimated using the command scan1coef and plotted using plot_coef in R/qtl2. The most likely genotypes of the analyzed F_2_ offspring at each QTL peak were extracted using the command maxmarg in R/qtl2. Phenotypes were plotted against genotypes at each QTL peak position and by combinations of all identified QTL peaks.

For each QTL region, we extracted all physical markers which mapped to genetic positions within these regions and identified all genes containing these physical markers. Gene lists were made using the annotation of the version 6 reference genome (Varadharajan et al. 2019) and liftover between version 6 and version 7 reference genomes (Rastas 2020; https://sourceforge.net/p/lep-anchor/code/ci/master/tree/liftover.awk).

## Results

### Genotyping and linkage maps

The final dataset consisted of 221 individuals which included four F_0_ grandparents, two F_1_ parents, and 215 F_2_ offspring (Fig. 1). The Lep-MAP3 Pileup2Likelihoods module identified 170 339 SNPs across the 21 linkage groups of which 84 526 informative sites were retained by ParentCall2. The largest 21 LGs identified by SeparateChromosomes2 contained 61 863 markers (Fig. S4) and the remaining 22 663 sites were designated as singletons. Finally, JoinSingles2All further added 7 485 sites into the linkage groups, resulting in a total of 69 348 sites separated into 21 LGs. Preliminary results showed that fewer than 8% of markers were paternally informative (i.e., using informativeMask=13) while more than 92% of the markers were maternally informative (informativeMask=23). This observation may not be surprising given that only the F_1_ mother was hybrid and that the males in general have fewer crossovers (Kivikoski et al. 2023). In addition, the *P. pungitius* reference genome was used which was more similar to the hybrid F_1_ female than the F_1_ male (*P. sinensis*). Therefore, only the maternally informative markers were utilized in the following analyses. The phased and matched maternal maps had a total of 60 089 sites, and 59 629 sites were retained after the manual cleaning (Fig. S5).

### QTL mapping of sex

The previous study by Natri et al. (2019) found that sex of hybrids between *P. sinensis* (ZW system) and *P. pungitius* (XY system in most lineages, unknown in WL) was controlled by the ZW system inherited from *P. sinensis* and mapped to a 19.4 cM region on LG12. In agreement with this, we found a strong association between phenotypic sex and LG12 and we narrowed down the sex determination region to two nearby QTL (in total 1 cM) with non-overlapping 95% CIs (Table 1). All F_2_ males were homozygous and F_2_ females heterozygous at the highest QTL peak at 11.826 cM except for 11 individuals (Fig. S6, S7) which were removed together with the individual of unknown sex (Table S1), retaining 203 individuals for the downstream analyses. The two QTL regions contained 43 SNPs located within six genes (*Usp7*, *EFS*, *DHRS4*, *Slc7a8*, *Svep1* and *TBC1D19*; Table S3).

**Table 1.** The QTL peaks for phenotypic traits. LG = linkage group, cM = centimorgan, PVE = percentage of variation explained.

| Phenotype | LG | Peak<br>(cM) | 95%CI<br>low<br>(cM) | 95%CI<br>high<br>(cM) | Range<br>(cM) | LOD | PVE<br>(%) | NO.<br>SNPs in<br>95% CI |
| --- | --- | --- | --- | --- | --- | --- | --- | --- |
| Sex | 12 | 11.286 | 10.819 | 11.286 | 0.467 | 61.08 | 74.98 | 18 |
| Sex | 12 | 13.623 | 13.623 | 14.090 | 0.467 | 51.25 | 68.74 | 25 |
| Standard Length | 12 | 87.896<sup>†</sup> | 81.591 | 92.927 | 11.336 | 5.35 | 11.43 | 139 |
| Standard Length | 16 | 43.088 | 32.301 | 45.438 | 13.137 | 7.21 | 15.08 | 110 |
| Body Height | 6 | 55.305 | 42.172 | 63.721 | 21.549 | 4.98 | 10.68 | 547 |
| Plate number | 4 | 137.699 | 133.021 | 137.699 | 4.678 | 4.58 | 9.87 | 1531 |
| Plate number | 13 | 64.174 | 60.436 | 65.109 | 4.673 | 9.09 | 18.64 | 428 |
| Plate number | 17 | 62.363 | 47.838 | 70.352 | 22.514 | 5.16 | 11.04 | 2677 |
| Plateness | 13 | 63.240 | 56.693 | 65.109 | 8.416 | 4.92 | 10.57 | 494 |
| Plate Area | 19 | 36.939<sup>‡</sup> | 30.427 | 58.042 | 27.615 | 3.68 | 8.01 | 2512 |
| Plate Area | 20 | 53.440 | 48.767 | 53.440 | 4.673 | 9.63 | 19.63 | 1642 |
| Plate Height | 20 | 53.440 | 49.234 | 53.440 | 4.206 | 13.02 | 25.58 | 1626 |
| Plate Width | 17 | 65.635 | 48.772 | 70.352 | 21.580 | 3.84 | 8.33 | 2663 |
| Plate Width | 19 | 33.707 | 4.677 | 56.172 | 51.495 | 3.26 | 7.12 | 2688 |
| Plate Width | 20 | 53.440 | 50.169 | 53.440 | 3.271 | 5.81 | 12.35 | 1597 |

<sup>†</sup> Pseudomarker (the nearest non-pseudo genetic position at 88.191).
<sup>‡</sup> Pseudomarker (the nearest non-pseudo genetic position at 36.043).

### QTL mapping of lateral plate phenotypes

Most of the lateral plate phenotypes were significantly dependent on phenotypic sex (Table S2). Females had more (x = 30.1 ± 0.45 [S.E.]) and bigger (x = 2.0 ± 0.10 mm^2^) lateral plates than males (plate number x = 28.7 ± 0.52, plate area: x = 1.6 ± 0.08 mm^2^). However, this sexual dimorphism is confounded by body size (females are larger, see below), as *t-*tests of plate number and plate size between sexes were not significant (*p* > 0.05) after controlling for standard length. Both plate number and the percentage of plated myomeres (*viz*. plateN_mean, plateness) were associated with a QTL at almost the same peak position in LG13 with overlapping confidence intervals (Table 1; Fig. 2A). In addition, plate number was associated with QTL on LG17 and LG4 (Fig. 2A), the latter containing the candidate gene *Eda* that controls lateral plate phenotypes in three-spined sticklebacks (Colosimo et al. 2004). Almost identical QTL regions were found when using the raw values of the plate number (plateN_mean; Table 1) and the transformed binary trait (plateN_binary; Table S4; Fig. S8), providing confidence of the identified regions being true QTL rather than false positives despite a skewed distribution of raw plate numbers. Both the LG4 and the LG13 QTL showed a positive effect of the *P. sinensi*s - *P. pungitius* hybrid genotype (SP) whereas the LG17 QTL showed a negative effect of the SP genotype on lateral plate numbers (Fig. 3A). Consistently, among the eight genotype combinations of these three QTL, the genotype having SS at both LG4 and LG13 but SP at LG17 generated the lowest number of lateral plates, while the reverse (SP at LG4 and LG13 while SS at LG17) generated the highest number of lateral plates (Fig. 3B).

**Figure 2.**
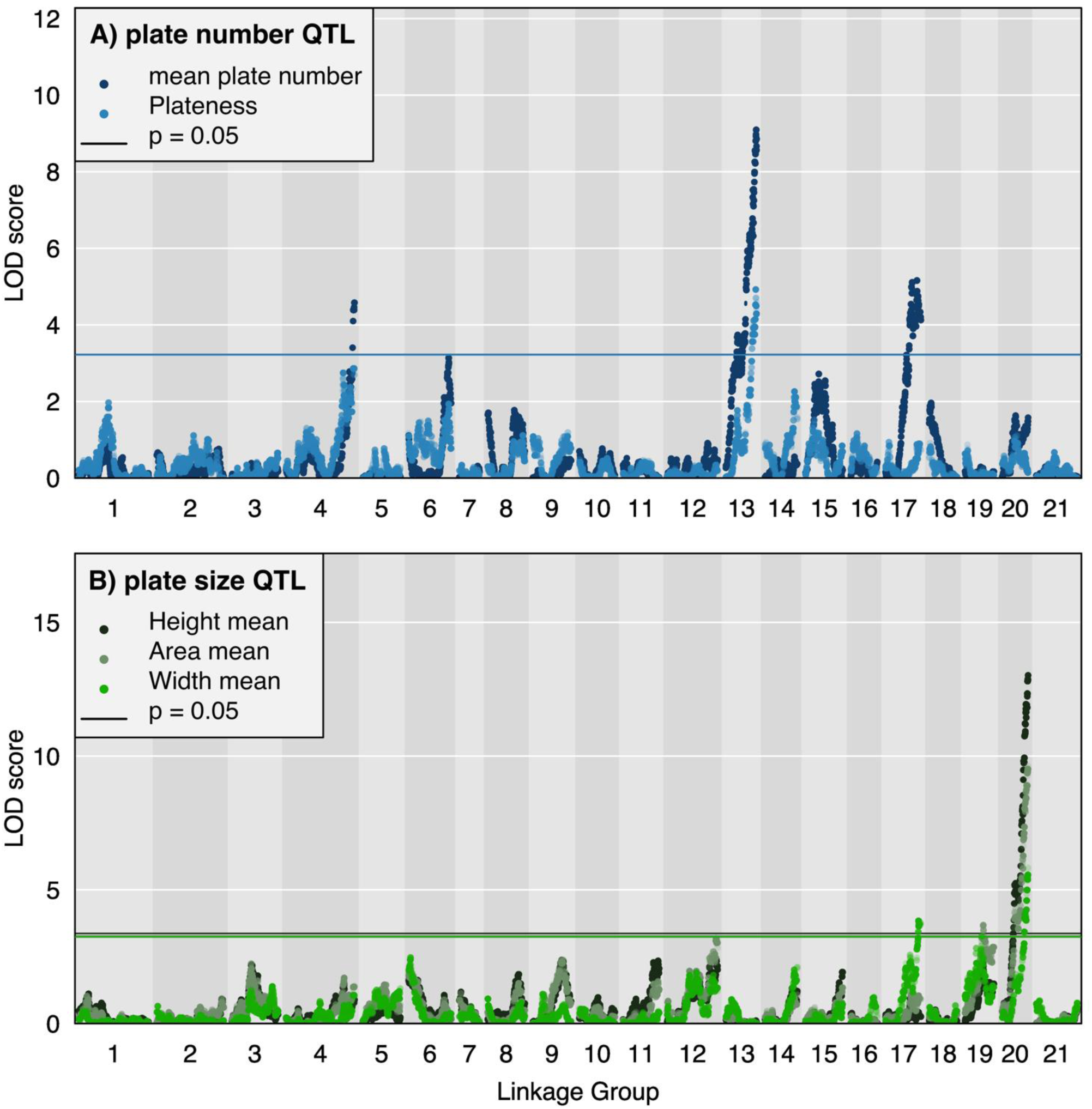
The QTL for lateral plate A) number and B) size in nine-spined sticklebacks. Horizontal lines show the significance threshold (*p* = 0.05) derived from 1000 permutation tests for each trait.

**Figure 3.**
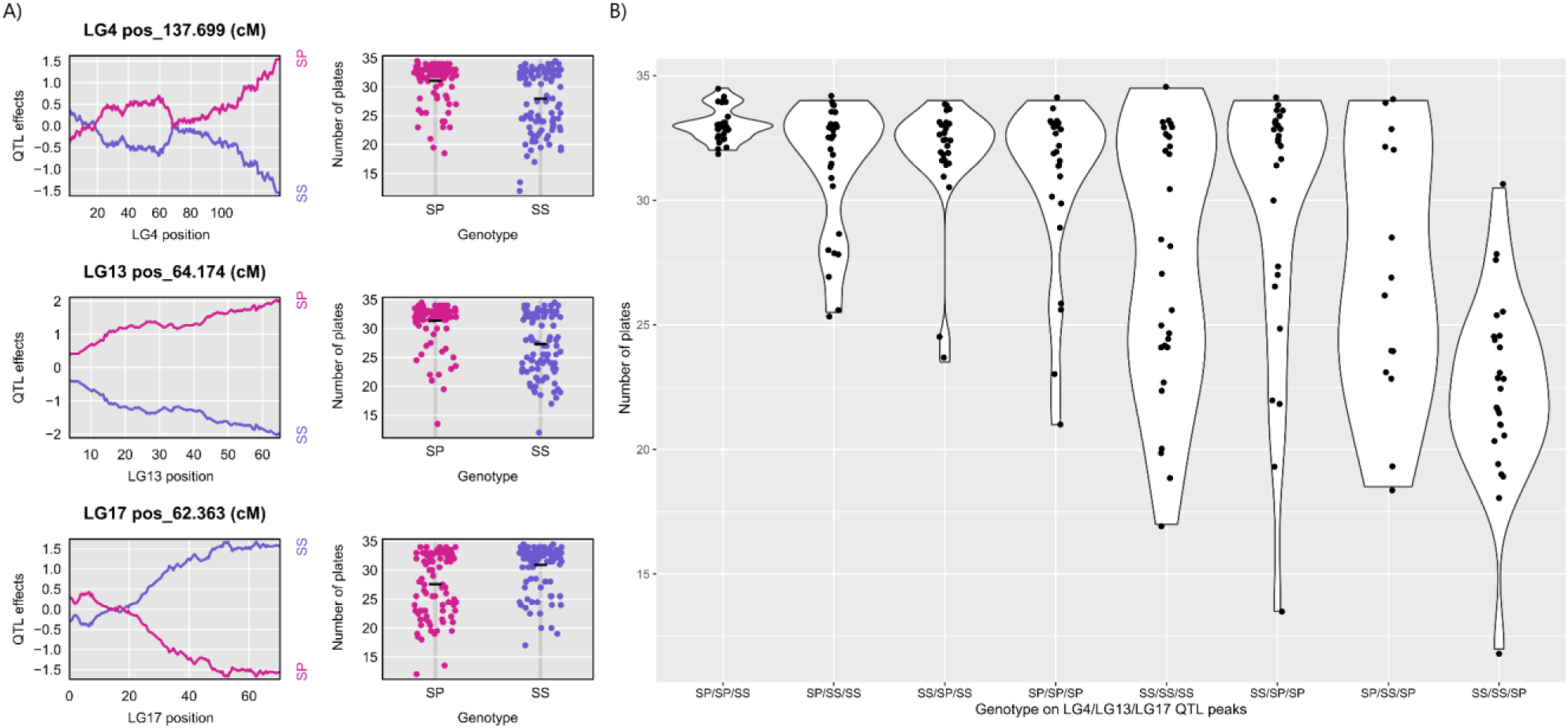
Effects of the genotypes at QTL peaks on lateral plate numbers. S represents the paternal allele from *P. sinensis* and P represents the maternal allele from *P. pungitius*. **A)** Effect size and phenotype-genotype plots of each QTL estimated by R/qtl2. **B)** Distribution of individual number of plates by genotype combinations at the three QTL.

The lasso regression approach produced a genomic estimate of heritability (*h^2^*) at 0.486 (Table S5), a value higher than the summed PVEs of the three plate number QTL (Sum_PVE_ = 0.396; Table 1), indicating that there might be additional QTL influencing plate number, albeit their contributions are likely to be minor.

The plate size phenotypes (*viz*. area, height, and width of the largest plate) were all associated with the same QTL peak on LG20 (Table 1; Fig. 2B). Plate area and plate width also mapped to nearby QTL peaks on LG19, but their 95% CIs were almost as wide as the entire LG including large regions of LOD scores below the significance threshold (Fig. S9). In addition, plate width also mapped to LG17, overlapping with the QTL for plate number (Table 1). All SP genotypes at these QTL peaks had positive effects except for the LG17 QTL in which SP had a negative effect on plate width (Fig. S10), similar to its negative effect on plate number (Fig. 3). Therefore, SP genotype at both LG19 and LG20 QTL peaks yielded the largest plate size and SP genotype at LG20 yielded the largest plate height, whereas SP genotype at both LG19 and LG20 and SS genotype at LG17 yielded the largest plate width (Fig. S11).

The lasso regression approach produced genomic heritabilities higher than the summed PVEs in plate area (*h^2^* = 0.495; Sum_PVE_ = 0.276), plate height (*h^2^*= 0.501; Sum_PVE_ = 0.256), and plate width (*h^2^* = 0.317; Sum_PVE_ = 0.278; Table 1, Table S5), suggesting that additional unidentified loci may influence variation in these traits.

### QTL mapping of body size

Females were significantly larger than males both in terms of standard length (SL; x_females_ = 52.9 ± 0.57 [S.E.] mm; x_males_ = 49.0 ± 0.44 mm; *t*_190.36_ = 5.33, *p* < 0.05) and body height (BH; x_females_= 11.0 ± 0.15 mm; x_males_= 10.4 ± 0.09 mm; *t*_174.57_ = 3.28, *p* < 0.05). Two moderate-effect QTL at LG12 and LG16 influenced SL and a QTL on LG6 influenced BH (Table 1; Fig. 4). The lasso method estimated genomic heritabilities of *h^2^* = 0.263 for SL and *h^2^* = 0.206 for BH (Table S5), and the corresponding summed PVEs across QTL were Sum_PVE_ = 0.265 for SL and Sum_PVE_ = 0.107 for BH (Table 1). It is noteworthy that the peak on LG6 in the analysis of SL was close to significance (Fig. 4), indicating that this region might also affect standard length by affecting the body height.

**Figure 4.**
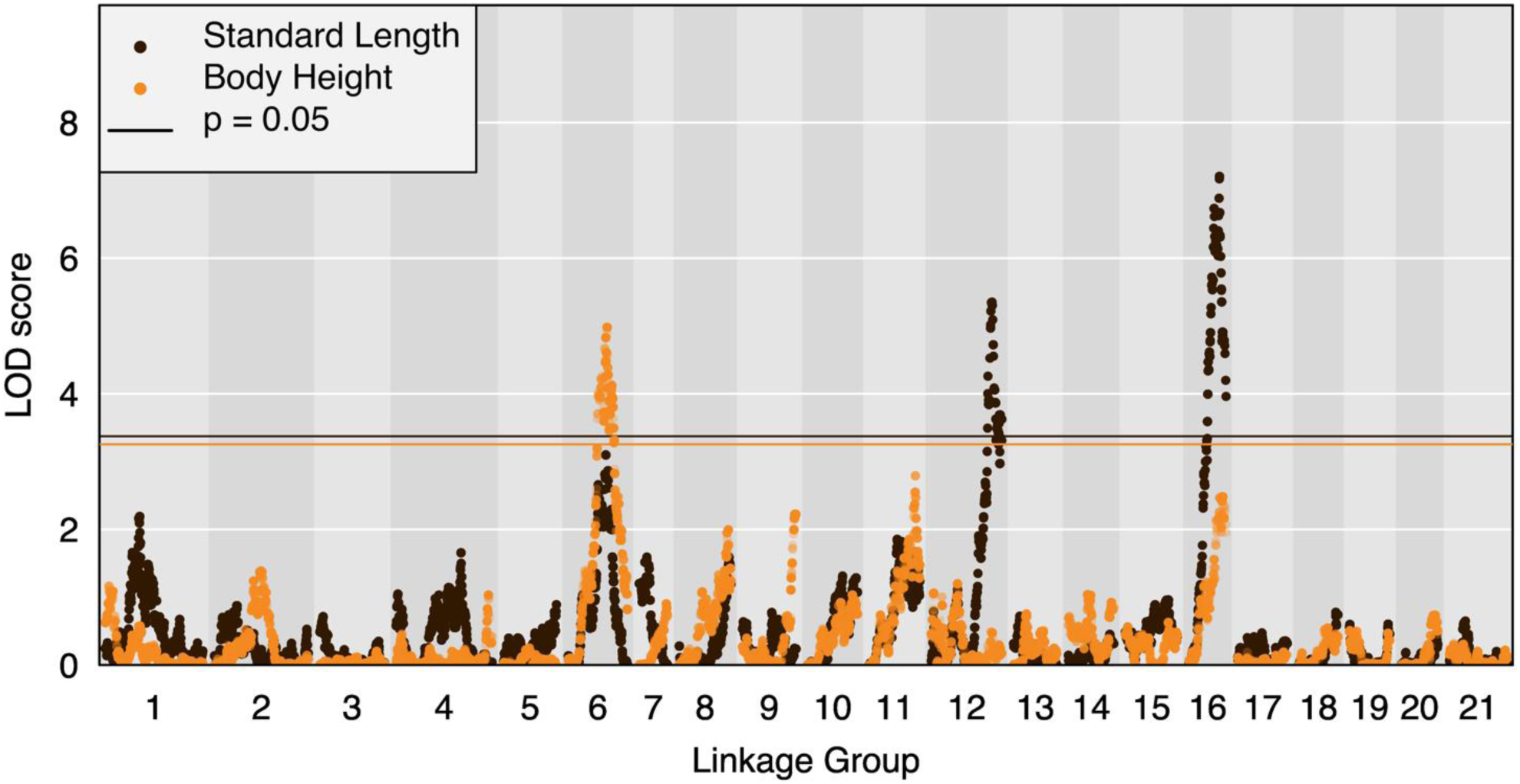
The QTL regions of body size variation. Horizontal lines show the significance threshold (*p* = 0.05) derived from 1000 permutation tests for each trait.

### Candidate genes

All genes containing SNPs in identified QTL regions are listed in Table S3. For the plate number variation, we identified several candidate genes whose functions might be associated with stickleback skeletal development, including the *Eda* gene on LG4, the fibroblast growth factors on LG4 (*Fgf4*, *Fgf16*, *Fgf20*, Jovelin et al. 2007), the bone morphogenetic protein genes on LG17 (*Bmp7*, Shawi et al. 2008; *Bmp11*, Indjeian et al. 2016), and potentially the BMP-2-inducible protein kinase on LG13 (*Bmp2k*, Li and Cao 2006; Wang et al. 2020). For the plate size variation, the *fgf23* on LG19 and the *Bmp7* and *Bmp11* on LG17 were identified as candidate genes. The BMP family in particular has been suggested to control the height and width of armor plates in sticklebacks (Indjeian et al. 2016). In addition, the paired box proteins (*Pax6* on LG19 and *Pax7* on LG17) have been found associated with the development of vertebrae in lab mice (Lang et al. 2007) and thus might also be candidate genes controlling the armor plate development in sticklebacks. The *Fgf14* on LG16 was identified as a candidate gene controlling the body size development.

## Discussion

Our study shows that variation in lateral plate number and plate size in the western European lineage (WL) of nine-spined sticklebacks were not fully controlled either by allelic variation in the same genetic loci as in the three-spined stickleback, or by the QTL controlling these phenotypes in the eastern European lineage (EL) of nine-spined sticklebacks. These results are consistent with the observation that pelvic armor morphology in the EL nine-spined sticklebacks is both more polygenic and heterogeneous (non-parallel) compared to the three-spined stickleback (Kemppainen et al. 2021). Hence, this study comprises yet another example of similar phenotypic changes being achieved by a variety of genetic architectures ranging from monogenic to oligo- or polygenic systems in the stickleback family Gasterosteidae (Colosimo et al. 2005; Shapiro et al. 2009; Yang et al. 2016; Kemppainen et al. 2021), rather than due to genetic parallelism governed by a few large-effect loci. Our findings highlight the notion that a diversity of genetic architectures can evolve to underpin similar phenotypic transitions, and that the frequency of reuse of the same genetic variation to achieve a given phenotypic transition (i.e., parallel evolution) seems to differ markedly between three- and nine-spined sticklebacks. Therefore, although freshwater populations of nine-spined sticklebacks are characterized by both small effective population sizes and low genetic connectivity (Shikano et al. 2010; Merilä 2014; Fang et al. 2021; Kivikoski et al. 2023), our results together with recent studies (Yang et al. 2016; Kemppainen et al. 2021) highlights that these two population demographic parameters alone do not necessarily limit evolutionary potential but rather result in a higher diversity of alleles underlying adaptive evolution.

Our results demonstrate that the genetic basis of lateral plate number variation in the WL nine-spined sticklebacks is controlled by at least three distinct moderate-effect QTL on different linkage groups, as well as unknown small-effect loci potentially across the genome as revealed by the lasso analyses. One of the moderate-effect QTL on LG4 contained the *Eda* gene known to be the major- effect locus typically explaining ~69% to 98% variation in lateral plate numbers in three-spined sticklebacks (Table 2; Colosimo et al. 2004; Liu et al. 2014; Glazer et al. 2015; Erickson et al. 2016; Schluter et al. 2021). However, this moderate-effect QTL explained only ~9.9% of variation in lateral plate numbers in WL nine-spined sticklebacks. The two other QTL associated with lateral plate number variation in our study were located on LG13 and LG17 and they accounted for 18.6% and 11.0% of variation, respectively. Hence, collectively the three QTL identified in this study accounted for up to 39.6% of variation in lateral plate numbers. Nearly similar amount of variation (33%) in this trait was explained by four QTL in the EL nine-spined sticklebacks (Table 2; Yang et al. 2016), but none of the QTL controlling lateral plate numbers are shared between EL and WL fish. Therefore, the genetic architecture of lateral plate numbers is divergent not only between the genera *Gastoresteous* and *Pungitius* but also within the genus *Pungitius*.

**Table 2.** Comparison of the genetic architecture underlying the variation of lateral plate number and size in stickleback fishes. Populations used in the cross experiments are given in parentheses. EL = eastern European lineage, WL = western European lineage.

| Species | Lateral plate number |  | Lateral plate size |  | Reference |
| --- | --- | --- | --- | --- | --- |
|  | QTL | PVE (%) | QTL | PVE (%) |  |
| *Gasterosteus aculeatus*<br>(Japan × Canada) | LG4 | 76.9 | LG4 | 11.5 - 12.9 | Colosimo et al.<br>2004 |
|  | LG7 | 3.7 | LG7 | 10.3 - 11.1 |  |
|  | LG10 | 6.5 | LG25<sup>†</sup> | 17.9 - 28.9 |  |
|  | LG26<sup>†</sup> | 4.5 |  |  |  |
| *Gasterosteus aculeatus*<br>(Finland) | LG4 | 74.4 | - | - | Liu et al. 2014 |
|  | LG9 | 9.1 |  |  |  |
|  | LG21 | 10.6 |  |  |  |
| *Gasterosteus aculeatus*<br>(Canada × US) | LG4 | 95.7 - 97.8 | - | - | Glazer et al.<br>2015 |
|  | LG7 | 13.3<sup>‡</sup> |  |  |  |
| *Gasterosteus aculeatus*<br>(Canada) | LG1 | 1.5 - 1.6 | - | - | Erickson et al.<br>2016 |
|  | LG3 | 2.4 |  |  |  |
|  | LG4 | 68.8 - 75.8 |  |  |  |
|  | LG5 | 4.5 |  |  |  |
|  | LG7 | 2.9 - 4.2 |  |  |  |
|  | LG21 | 1.6 - 7.5 |  |  |  |
| *Gasterosteus aculeatus*<br>(Canada) | LG4 | 85.8 | - | - | Schluter et al.<br>2021 |
| *Pungitius pungitius*<br>(Canada × Alaska) | LG12<sup>§</sup> | 28.4 - 30.1 | - | - | Shapiro et al.<br>2009 |
| *Pungitius pungitius*<br>(Finland EL) | LG8 | 8.7 | - | - | Yang et al.<br>2016 |
|  | LG12 | 7.3 |  |  |  |
|  | LG20 | 8.0 - 11.3 |  |  |  |
|  | LG21 | 8.6 - 10.0 |  |  |  |
| *Pungitius pungitius*<br>(Japan *P. sinensis* ×<br>Belgium WL) | LG4 | 9.9 | LG17 | 8.3 | This study |
|  | LG13 | 18.6 | LG19 | 7.1 - 8.0 |  |
|  | LG17 | 11.0 | LG20 | 12.4 - 25.6 |  |

<sup>†</sup> The original linkage group studies in *Gasterosteus aculeatus* were based on a linkage map with 26 LGs before consolidation to the 21 confirmed chromosomes. <sup>‡</sup> Residual variance in *Eda* heterozygotes <sup>§</sup> The same region control sex

Similar to the results regarding genetic architectures of later plate number variation, the three QTL controlling lateral plate size variation in WL nine-spined sticklebacks were different from those in three-spined sticklebacks (Table 2). These three QTL regions (on LG17, LG19, and LG20) accounted for up to 28% of variation in the lateral plate size. Because large effect sizes are more likely to be significant and detected as QTL regions, genetic loci of small effect sizes may not be detected due to lack of statistical power. In accordance with this, estimates from lasso regressions suggested that there are yet undetected small-effect loci (among the non-QTL SNPs) accounting for the unexplained variation (37% in plate area, 34% in plate height, and 20% in plate width, Table S5), indicating oligo- to polygenic inheritance of these traits. These results are consistent with the previous study showing that genetic architectures of pelvic reduction in nine-spined sticklebacks range from monogenic to oligogenic and polygenic inheritance (Kemppainen et al. 2021).

Among the moderate-effect QTL (on LG6, LG12, and LG16) explaining variation in standard length and body height of the WL nine-spined sticklebacks, only the LG12 QTL is shared with the QTL affecting body size in the EL nine-spined sticklebacks (on LG8, LG12, and LG13; Laine et al. 2013). Similar to the results regarding lateral plate variation, the body size variation also appears to have different genetic architectures between EL and WL fishes. However, it is unclear whether the two LG12 QTL regions are actually the same or not in EL and WL, and it is hard to judge how different their genetic architectures are because our QTL analyses likely did not have power to detect loci of small effects. Nevertheless, genomic heritabilities for standard length and body depth (*h^2^* = 0.21 - 0.26) were very similar to the summed PVEs of individual QTL affecting these traits (0.11 - 0.27), suggesting an oligogenic basis of body size variation. This inference is supported by biometric estimates which have yielded very similar heritabilities within (*h^2^* = 0.13 - 0.18; Shimada et al. 2011) and among (*h^2^* = 0.07 - 0.25; Fraimout et al. 2022) population crosses in nine-spined sticklebacks.

We confirmed findings in Natri et al. (2019) that the ZW system inherited from *P. sinensis* controlled phenotypic sex of the hybrids between *P. sinensis* and *P. pungitius*. Our study also narrowed down the sex determination region of *P*. *sinensis* from around 19.4 cM in Natri et al. (2019) to two non-overlapping QTL regions of around total 1 cM on LG12 which explained 69% - 75% of the variation in sex. Recent studies found a translocated copy of the anti-Müllerian hormone gene (*Amh*) on the Y chromosome (*Amhy*) to be the candidate sex determination gene in several stickleback lineages (Jeffries et al. 2022), including the three-spined stickleback (Peichel et al. 2020), the blackspotted stickleback (*Gasterosteus wheatlandi*, Sardell et al. 2021), and the brook stickleback (*Culaea inconstans*, Jeffries et al. 2022), the latter being the sister genus of *Pungitius* (Guo et al. 2019). However, only the ancestral copy of the *Amh* gene is annotated in the nine-spined stickleback reference genome (on LG8; Varadharajan et al. 2019), indicating that it is unlikely the sex determination gene in *Pungitius* sticklebacks. In fact, so far no sex determination gene has been identified in any *Pungitius* sticklebacks despite known sex chromosomes (Jeffries et al. 2022). In this study, the SNPs mapped to sex-associated QTL regions fell into six annotated genes, among which the Ubiquitin carboxyl-terminal hydrolase 7 (*Usp7*) has been found to regulate sex chromosome development and interact with the sex determination region in mice (Luo et al. 2015; Dong et al. 2021). Therefore, *Usp7* is a candidate gene for sex determination of *P. sinensis*.

Among the genes that have been previously identified to have strong associations with the three-spined stickleback morphological development (summarized in Reid et al. 2021), only *Eda* on LG4 was included in our candidate gene list for the lateral plate number. Furthermore, we identified *Bmp2k* on LG13 as a candidate gene that impacts plate number variation in the nine-spined stickleback, and *Bmp7* on LG17 as a gene that potentially influences both plate number and plate width (although not plate area or height). Similarly, a recent study in the three-spined stickleback also found that the BMP pathway is a strong candidate mediating the effect of *Eda* on the development of lateral plates (Rodríguez-Ramírez et al. 2023). However, as all of the QTL regions in our study included tens to hundreds of genes, little confidence can be assigned to the identified candidate genes. Further narrowing of the QTL regions would require breeding individuals beyond F_2_ generation, or adopting genome-wide association approaches with larger sample sizes than what was available for this study.

The strong isolation by distance in the sea and severe bottlenecks associated with freshwater colonisations have led to the prediction that nine-spined sticklebacks are more limited in reaching their phenotypic optima when colonising freshwater habitats compared to three spined sticklebacks (Fang et al. 2021). This prediction was supported by the observation that in three freshwater (pond) populations of EL nine-spined sticklebacks, pelvic reduction was only complete in one population where it was controlled by a single large-effect locus (*pitx*1), whereas the other two populations only showed some level of pelvic reduction possibly controlled by alternative medium- or small-effect alleles at other QTL regions (Kemppainen et al. 2021). Similarly, here we showed that the reduction of lateral plate numbers in freshwater WL nine-spined sticklebacks is controlled by multiple medium- to small-effect loci that are different from those in EL nine-spined sticklebacks and the large-effect *Eda* gene in three-spined sticklebacks. Accordingly, empirical evidence rejected the hypothesis of parallel evolution in the repeatable phenotypic changes of freshwater *Pungitius* sticklebacks, but indicated non-parallel and possibly redundant genetic architectures leading to similar phenotypic changes. Such redundancy is likely crucial for the evolutionary potential of species and populations facing rapid environmental changes, especially in fragmented habitats causing increased genetic differentiation and bottlenecks. For the species with fragmented small populations, redundant genetic architectures increase the likelihood that at least some populations can adapt to some novel environmental pressures, but it is difficult to achieve adaptation in all traits within one population, and the long term species survival probability will still depend on the spread of adaptive alleles across populations via gene flow.

## Conclusions and outlook

In conclusion, our results add to the growing evidence (Shapiro et al. 2009; Shikano et al. 2013; Yang et al. 2016; Fang et al. 2020; Kemppainen et al. 2021) that the repeatability of evolutionary transitions in response to similar selection pressures in *Pungitius* sticklebacks is often underlied by different genetic architectures of moderate- to small-effect loci. This is in stark contrast to the situation in the closely related three-spined stickleback which tends to utilize the same major-effect loci in independent colonizations of freshwater habitats. However, even in the case of the three-spined stickleback, growing evidence suggests that genetic parallelism in adaptation to freshwater is not as prevalent as early studies conducted in the Eastern Pacific region suggested: a greatly reduced degree of genetic parallelism has been found in the Atlantic basin likely due to variation lost during colonizations as well as reduced gene flow among these populations (DeFaveri et al. 2011; Fang et al. 2020; Dahms et al. 2022). The same also applies to stream-lake populations of three-spined sticklebacks experiencing reduced gene flow (Conte et al., 2012; Conte et al., 2015; Stuart et al., 2017; reviewed in Kemppainen et al. 2021). We suggest that further focus on genetic underpinnings of phenotypic variation in the morphologically and genetically diverse stickleback genus *Pungitius* (Guo et al. 2019) can provide a useful framework to quantify how the likelihood of genetic parallelism underlying similar phenotypic changes decreases with increasing evolutionary distance between taxa (Elmer and Meyer 2011; Conte *et al*. 2012) and also inform us about variation in species’ responses to changing environments (Bay et al. 2017).

