## Supplementary Materials for "Heterogeneous genomic architecture of skeletal armour traits in sticklebacks"

### Supplementary information

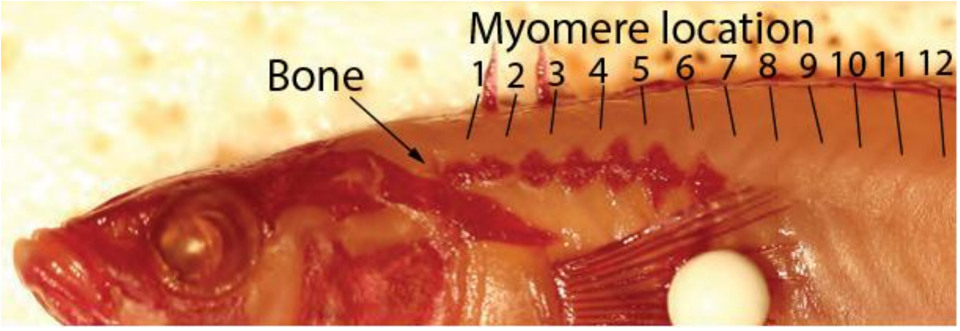


**Figure S1. An example of the myomere positions in a stained fish.**

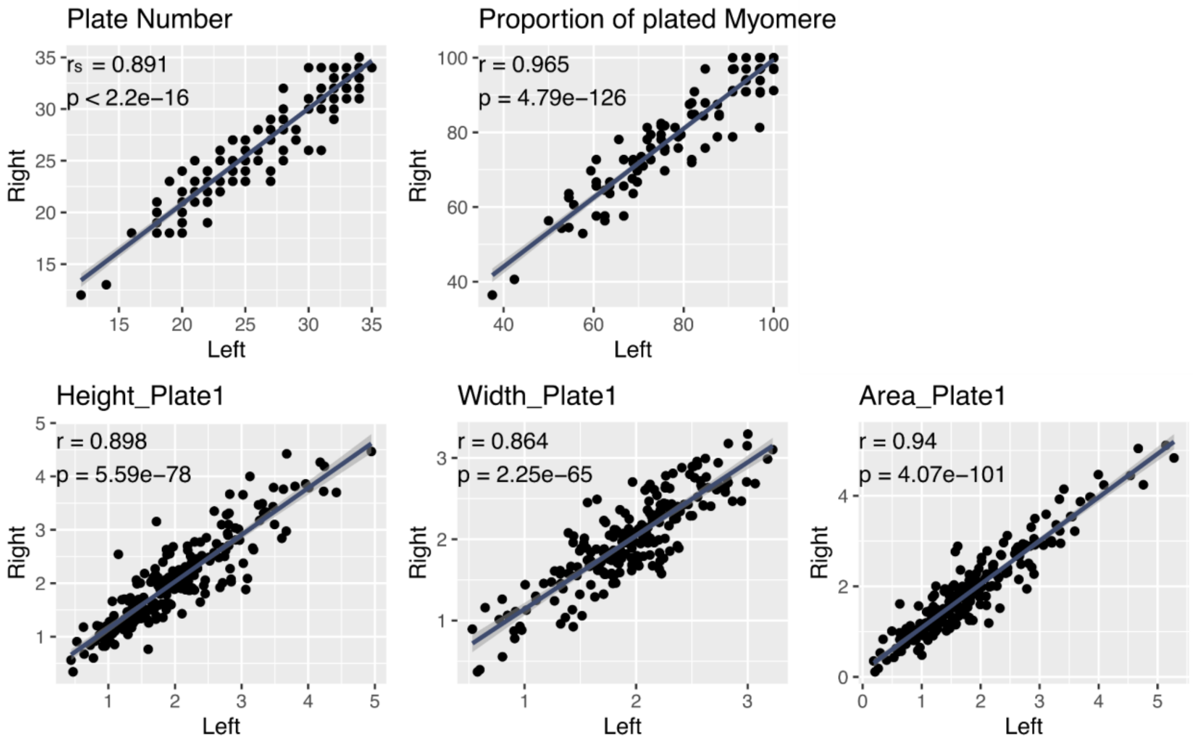


**Figure S2. Correlation of the measurements on the left and right side lateral plate traits.** Regression lines were added by fitting data to the linear model for illustrative pursposes. Plate 1 refers to the largest plate on either side.

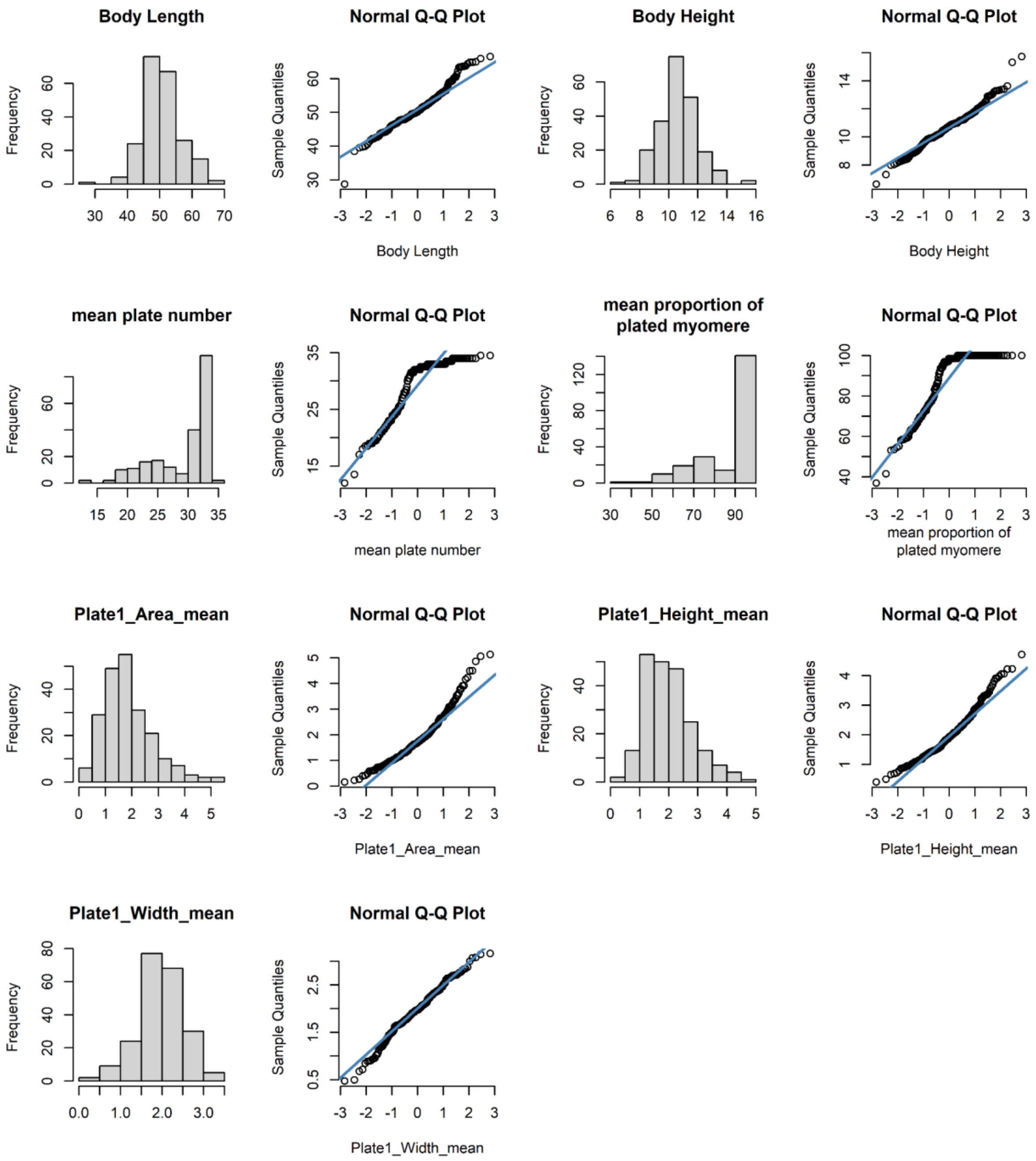


**Figure S3. Distribution of the phenotypic measurements.** Most traits conferred to a normal distribution. Plate 1 refers to the largest plate on either side. Plate area has a very slight skew to the right but log transformation did not improve this (not shown). Plate number and the proportion of plated myomeres were strongly skewed, and they were transformed into binary traits in the QTL analyses (see main text) .

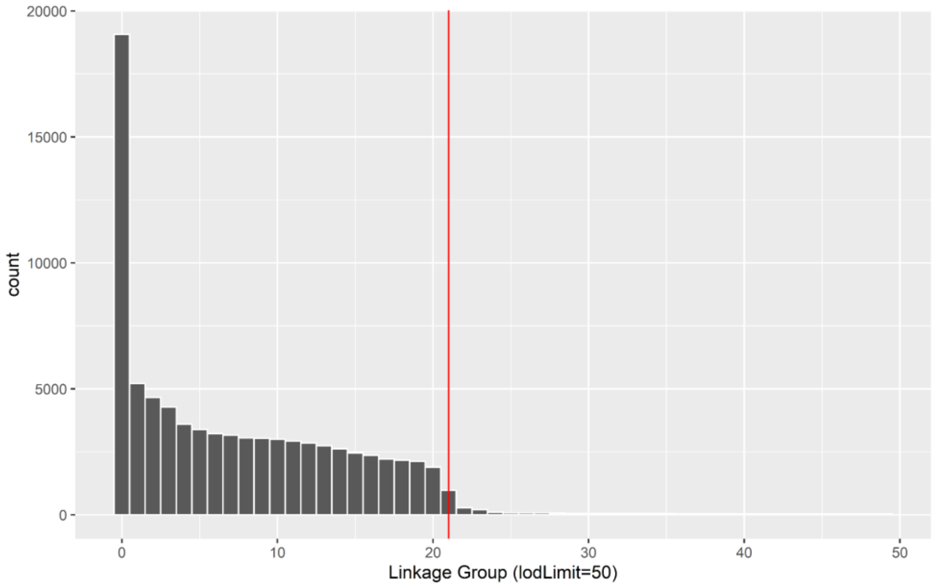


**Figure S4. The first 50 linkage groups generated from SeparateChromosomes2.** Y-axis represents the number of SNPs assigned to each linkage group (LG). LGs are ordered by the number of SNPs and the LG 0 represents singletons (SNPs not assigned to any linkage group). The red line highlights LG 21 and the subsequent LGs were merged with LG 0 (see main text).

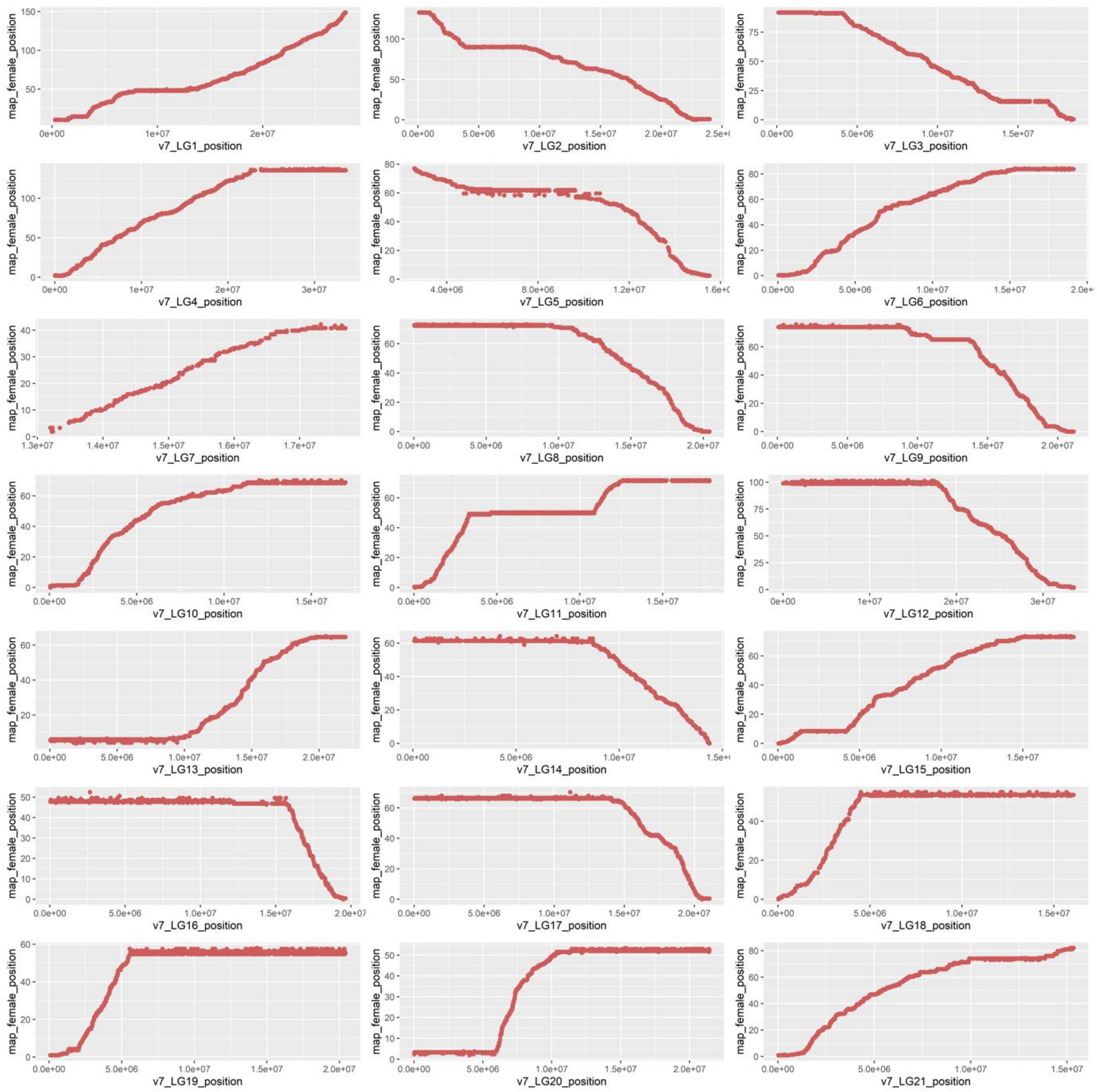


**Figure S5. Marey plots of the manually cleaned linkage maps.** The x axis represents physical positions on each linkage group of the version 7 reference genome. The y axis represents genetic positions in the linkage maps. Only female positions were plotted because we only used the maternal-informative sites (see main text) and thus no male patterns were retained. The flat regions indicated inversions between the two crossed species (*Pungitius sinensis* and *P. pungitius*).

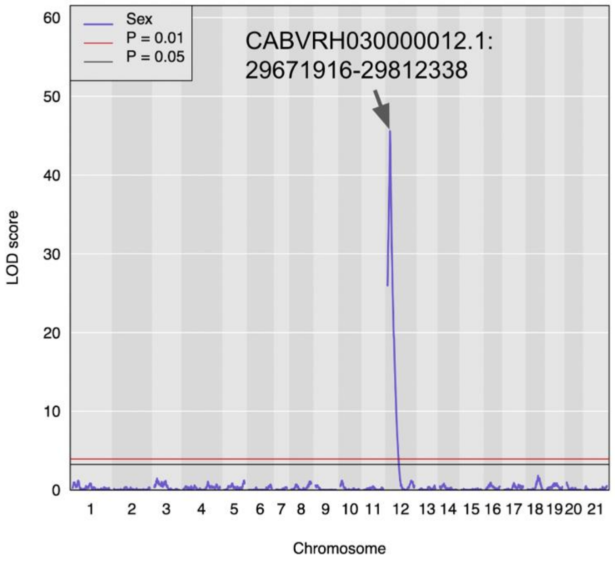


**Figure S6. QTL mapping of sex using all 215 individuals.** Horizontal lines show the significance thresholds (*p*=0.05 and *p*=0.01) derived from 1000 permutation tests. One peak (LOD=45.58) was identified on LG12 at the genetic position 11.286 cM (95% CI 10.819 - 11.754). The physical markers mapping to this peak included 15 SNPs located between 29671916 – 29812338 bp on LG12 (labelled in the figure). All phenotypic males had homozygous genotypes at these SNPs except for the individuals C1-F15, C1-F18, C1-F19, C3-1, C3-29, and C3-7. All phenotypic females had heterozygous genotypes at these SNPs except for the individuals C1-F9, C2-37, C3-23, C3-34, and C7-8. Therefore, these 11 individuals together with an individual of unknown phenotypic sex were removed from downstream analyses.

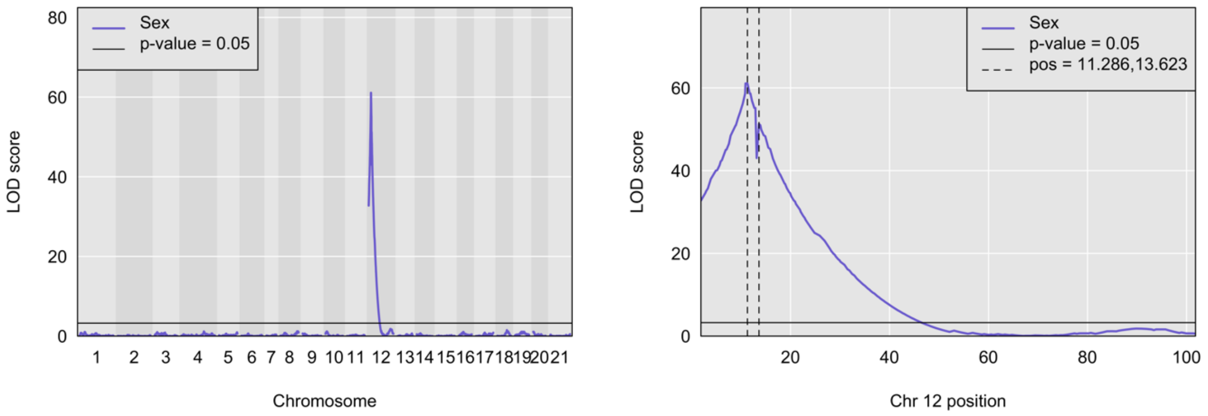


**Figure S7. QTL mapping of sex using the remaining 203 individuals.** Horizontal lines show the significance threshold (*p*=0.05) derived from 1000 permutation tests. A higher maximum number of permutation iterations (maxit=120) was used to achieve regression convergence. Two peaks were identified on the LG12 and their positions (cM) were labelled on the right plot. Among the 203 individuals, all females were heterozygous and males homozygous at the peak at 11.286 cM (15 physical SNPs), but there was still one homozygous female and four heterozygous males at the peak at 13.623 cM (8 physical SNPs).

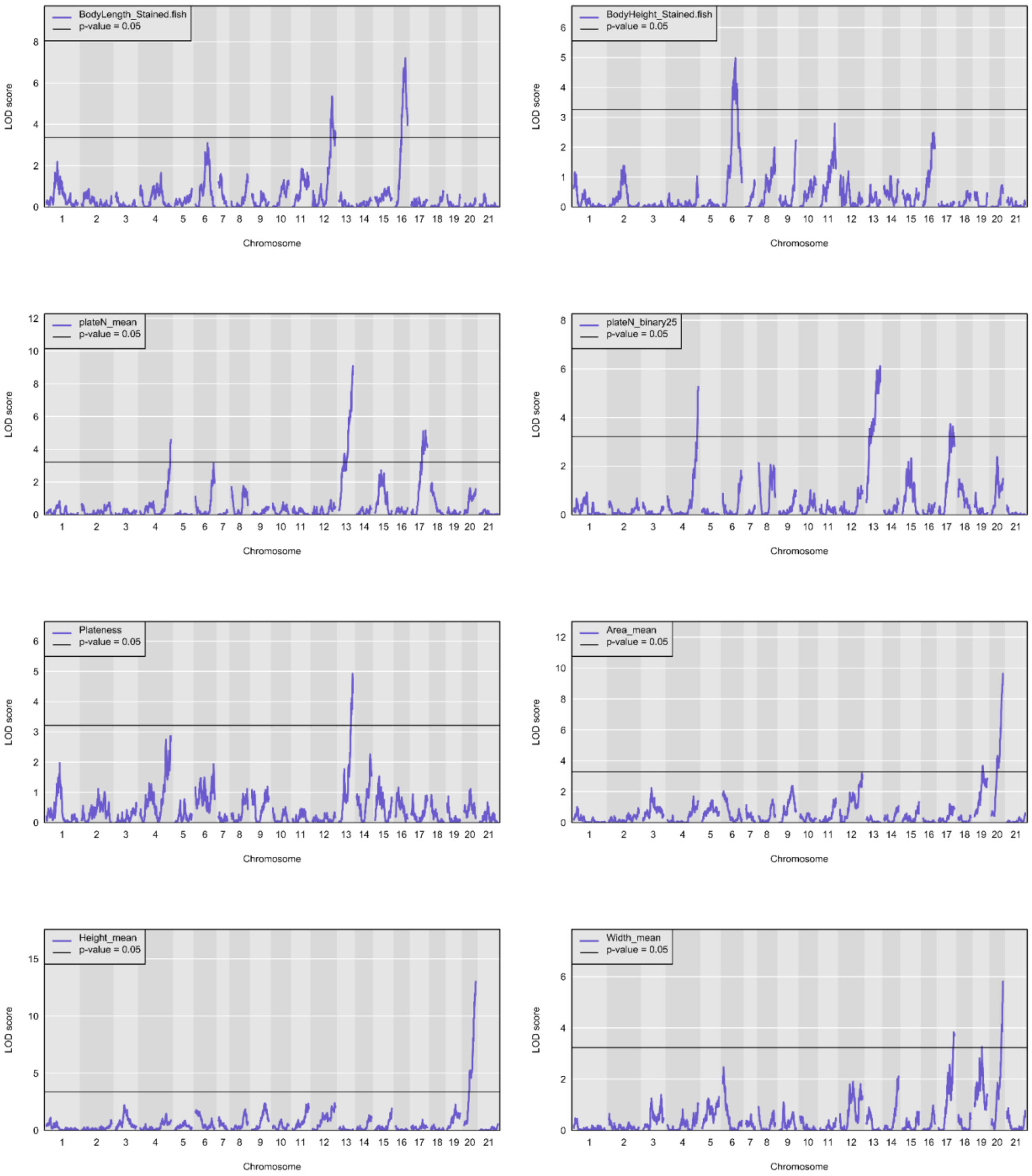


**Figure S8. QTL mapping of each phenotype.** Horizontal lines show the significance threshold (*p*=0.05) derived from 1000 permutation tests.

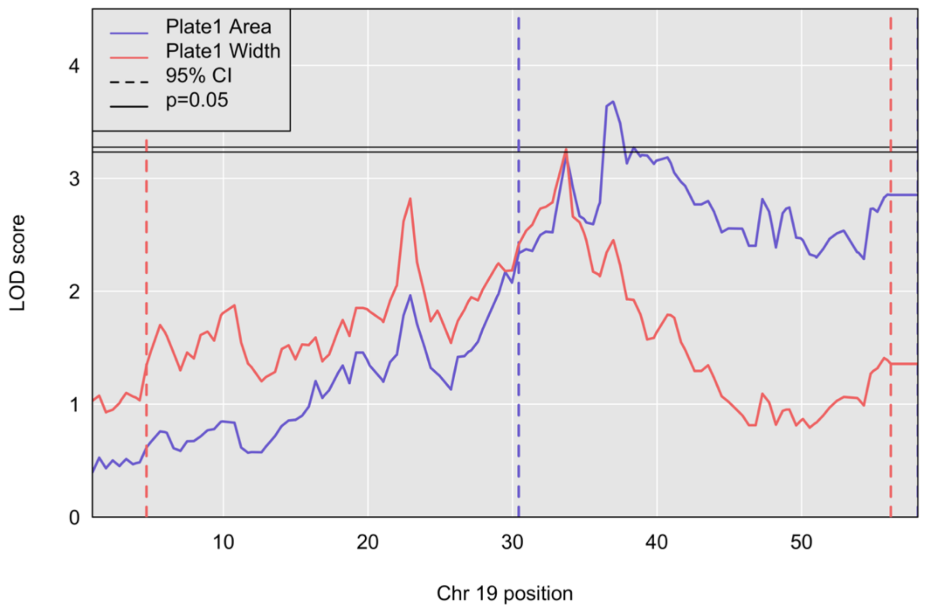


**Figure S9. The QTL region on LG19 mapped to the plate area and plate width.** Horizontal lines show the significance threshold (*p*=0.05) derived from 1000 permutation tests for each trait. Vertical dashed lines show the range of the 95% CI.

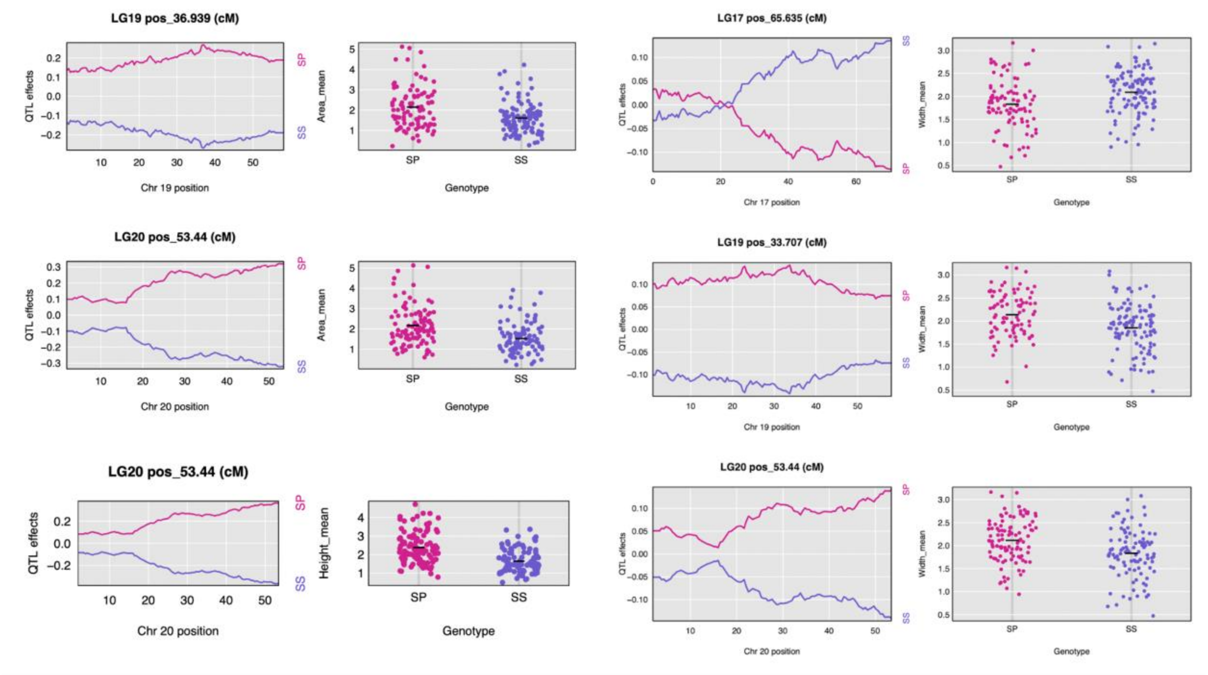


**Figure S10. Effect size and phenotype-genotype plots of each QTL peak for the plate size variation.** S represents the paternal allele from *P. sinensis* and P represents the maternal allele from *P. pungitius*.

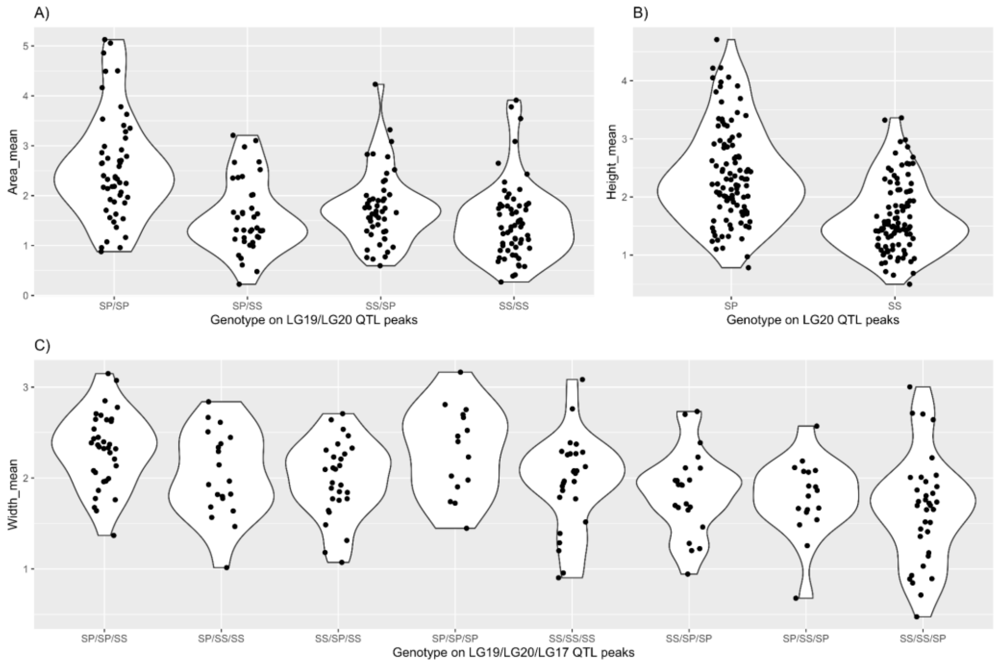


**Figure S11. Plate size variation and their genotypes at corresponding QTL peaks.** S represents the paternal allele from *P. sinenesis* and P represents the maternal allele from *P. pungitius*. **A)** The mean area of the largest plate on both sides. QTL regions were found on LG19 and LG20. **B)** The mean height of the largest plate on both sides. QTL regions were found on LG20. **C)** The mean width of the largest plate on both sides. QTL regions were found on LG19, LG20, and LG17.

**Table S1. Sample information and phenotypic measurements.** The dataset listed all 221 samples including the four $F_0$ grandparents, two $F_1$ parents, and 215 $F_2$ offspring. Raw phenotypic data of all $F_2$ individuals are included. The 12 individuals with potentially misidentified or unknown phenotypic sexes were marked.

*Note: Table S1 is attached as a separate excel.*

**Table S2. Dependency tests of the numeric phenotypes.** Significance levels were indicated by asterisks (* $p$<0.05; ** $p$<0.01; *** $p$<0.001). The significant independent variables (in bold) were added as additive covariates in the QTL mapping of the corresponding phenotype.

| | ANOVA | | | Linear model on the residuals | | |
|---|---|---|---|---|---|---|
| dependent variable (phenotype) | independent variable | *F* | *p* | independent variable | *t* | *p* |
| Standard Length | **Sex** | 29.74 | 1.45e-07 *** | - | - | - |
| | **Clutch** | 3.506 | 0.00466 ** | - | - | - |
| Body Height | **Sex** | 11.088 | 0.00104 ** | - | - | - |
| | **Clutch** | 3.255 | 0.00759 ** | - | - | - |
| plateN_mean | **Sex** | 3.987 | 0.0472 * * | Standard Length | 0.422 | 0.673 |
| | Clutch | 0.648 | 0.6634 | Body Height | -0.882 | 0.379 |
| Plate1 Area | **Sex** | 9.721 | 0.0021 ** | **Standard Length** | 5.763 | 3.08e-08 *** |
| | Clutch | 1.303 | 0.2644 | Body Height | -1.123 | 0.263 |
| Plate1 Height | Sex | 1.189 | 0.2768 | **Standard Length** | 4.174 | 4.46e-05 *** |
| | Clutch | 2.237 | 0.0521 | Body Height | -0.874 | 0.383 |
| Plate1 Width | **Sex** | 20.16 | 1.22e-05 *** | **Standard Length** | 5.274 | 3.46e-07 *** |
| | **Clutch** | 2.46 | 0.0345 * | Body Height | -0.696 | 0.487 |

**Table S3. List of all genes that include any physical sites mapped to the QTL regions.** Phenotype were listed on separate sheets.

*Note: Table S3 is attached as a separate excel.*

**Table S4. The QTL peaks for the transformed trait plateN_binary.**

| Phenotype | LG | Peak (cM) | 95%CI low (cM) | 95%CI high (cM) | Range (cM) | LOD | PVE (%) | # physical sites in 95% CI |
|---|---|---|---|---|---|---|---|---|
| plateN_binary | 13 | 64.174 | 45.469 | 65.109 | 19.640 | 6.12 | 12.97 | 823 |
| plateN_binary | 4 | 137.699 | 133.021 | 137.699 | 4.678 | 5.26 | 11.26 | 1531 |
| plateN_binary | 17 | 51.580 † | 47.838 | 70.352 | 22.514 | 3.73 | 8.11 | 2677 |

† Pseudomarker (the nearest non-pseudo genetic position at 52.00).

**Table S5. Heritability of each trait estimated using the lasso regression.** All 2 677 SNPs were used to estimate the total PVE which approximates the narrow sense heritability.

| Trait | PVE.total (heritability) | SNPs | PVE.chro (%) |
|---|---|---|---|
| Standard Length | 0.2627 | nonQTL | 5.52 |
| | | LG12 | 7.73 |
| | | LG16 | 12.53 |
| Body Height | 0.2064 | nonQTL | 13.52 |
| | | LG6 | 8.51 |
| Mean plate number | 0.4855 | nonQTL | 14.34 |
| | | LG4 | 7.09 |
| | | LG13 | 18.24 |
| | | LG17 | 9.26 |
| Plateness | 0.2936 | nonQTL | 20.59 |
| | | LG13 | 9.09 |
| Plate1 Area | 0.4953 | nonQTL | 37.49 |
| | | LG19 | 3.86 |
| | | LG20 | 6.67 |
| Plate1 Height | 0.5006 | nonQTL | 33.57 |
| | | LG20 | 15.04 |
| Plate1 Width | 0.3172 | nonQTL | 20.25 |
| | | LG17 | 3.16 |
| | | LG19 | 3.65 |
| | | LG20 | 5.45 |
